# Can *Colpoda* travel across oceans? Salinity tolerance of resting cysts may enable global dispersal of the species

**DOI:** 10.1101/2023.11.22.568208

**Authors:** Ryota Saito, Hiroki Yamanobe, Kazuma Yabuki, Tomohiro Suzuki, Takeru Saito, Shuntaro Hakozaki, Manfred Wanner, Ryota Koizumi, Tatsuya Sakai, Maribet Gamboa, Toshihiko Tanaka, Akiko Ono, Hoa Thanh Nguyen, Yuta Saito, Tetsuya Aoyama, Katsuhiko Kojima, Futoshi Suizu, Kozo Watanabe, Yoichiro Sogame

## Abstract

Protist species are distributed worldwide. The processes that enabled this global distribution are unclear. One possible means is through the oceanic dispersal of freshwater protists, although this has not been investigated in detail to date. Here, the ability of resting cysts to tolerate saline conditions is examined as a possible mechanism that enables the oceanic dispersal of protists. Resting cysts of *Colpoda cucullus*, a freshwater soil protist, can tolerate saline conditions of at least 3.5% NaCl for more than one week. A transcriptome analysis showed that the relative levels of expression of genes associated with membrane function are increased in resting cysts, indicating that salinity tolerance is associated with reconstruction of the cell membrane. Additionally, the outer layer of the cyst wall, a shell-like ectocyst, includes chitin. This may function as a form of “biological armor” that protects the cell from physical stress during oceanic dispersal.

Teaser: Fresh water protists may be dispersed across oceans by formation of resting cysts that can tolerate saline conditions.

## Introduction

Dispersal and biodiversity of species has fascinated scientists since the times of Carl von Linné and Charles Darwin. Unicellular protists, such as ciliates, have survived drastic environmental changes over more than a billion years since eukaryotic life emerged (1) without evolving into multicellular organisms. Hence, they offer a means to investigate the fundamental aspects of dispersal and biodiversity.

With regard to unicellular organisms, the theoretical idea of ‘everything is everywhere, but the environment selects’ is often used to describe biodiversity (2–5). Protist species have been stated to be ubiquitous (6) and are present in any location that has a suitable habitat; their widespread distribution has led to protists being described as ’cosmopolitan’ species (7). Each local species is rich and hence its global richness is represented by the diversity of the local species (6, 8-10). Globally-distributed species are generally described as rich, while “flagship” species may have distinctive morphologies and show endemic and restricted biogeographical distribution (11,12). There is still considerable debate on the differences between cosmopolitan and flagship species. It is known that one-third of protist species have a restricted distribution, whereas others are thought to be cosmopolitan (13). Additionally, the use of molecular data has provided insights into ‘cryptic diversity’ in many organisms such as testate amoebae (13–18) and marine protists (19). Thus, supposedly cosmopolitan species may be in fact be a “hidden” or cryptic complex of different species with some geographical restrictions (20,21). Can cosmopolitan species achieve global diversity ?

In protists, global diversity is a result of dispersal due to geological history, human introduction, and transportation by wind and animals in the cystic state (12,13). Another possible route of dispersion is via oceans when in the resting cyst stage. This study examines this possibility.

Encystment via resting cysts is the strategy used by protists to survive in unfavorable environments (22–24). In cysts, metabolic activity is reduced to an unmeasurable level; reversal of encystment occurs through a process of excystment when conditions become more favorable (22, 25, 26). During encystment in *Colpoda*, the cell is surrounded by a cyst wall formed by an outermost layer (the ectocyst) and several inner layers (the endocyst) (27,28). The cysts have little metabolic activity but have extreme tolerance against environmental stresses such as desiccation (29), high and low temperatures (30), freezing (31,32), UV irradiation (33), acids and alkalis (34,35), electrostatic exposure (36), and gamma irradiation (37,38). Although water is an essential factor in protist survival, they can nevertheless survive even in arid terrestrial environments by forming resting cysts. However, saline water is inimical to the survival of freshwater soil protists such as *Colpoda*.

Can *Colpoda* be dispersed across oceans? They may be able to do so by forming resting cysts. However, to date, there are no reports on salinity tolerance of *Colpoda* cysts. For oceanic survival, the cysts must also protect against physical damage in addition to resisting the deleterious effects of saline conditions. In the present study, we investigated the ability of cysts to tolerate saline conditions and resist physical damage. The data should provide new insights into the global dispersal of freshwater protists.

## Materials and Methods

### Culture, induction of encystment and excystment, sample preparation

*Colpoda cucullus* strain R2TTYS was maintained in culture medium (infusion of 0.05% (w/v) dried rice leaves supplemented with 0.05% Na_2_HPO_4_). After 1 day of culture, cells were collected by centrifugation (1500 x g for 1 min), washed twice in 1 mM Tris-HCl (pH 7.2), and used for experiments. Encystment was induced by suspending *C. cucullus* cells at high cell density (10,000–50,000 cells mL^-1^) in encystment-inducing medium (1 mM Tris-HCl (pH 7.2), 0.1 mM CaCl_2_; En-medium). For RNA extraction, 50 μg/mL (final conc.) ampicillin sodium was added to the samples. Excystment was induced by replacing En-medium with excystment-inducing medium (0.2% infusion of dried rice leaves supplemented with 0.05% (w/v) Na_2_HPO_4_; Ex-medium).

### Microscopy

Vegetative cells and cysts of *C. cucullus* were analyzed under an optical microscope Axiovert A1 system (Carl Zeiss Co. Ltd., Tokyo, Japan) or Axioscope A1 system (Carl Zeiss Co. Ltd.). Endocyst staining was performed using toluidine blue (TB) as previously described (34).

The effects of salinity on the cysts were assessed using a “shrink ratio” as described previously (35). In brief, images of 1-week-old cysts (Fig. 2A‘C’) exposed to 3.5 % NaCl for a week were captured. Micrographs from the prepared cysts (Fig. 2A‘P’) were taken, washed with 0.1 mM CaCl_2_/1 mM Tris-HCl (pH 7.2), and micrographs were taken again (Fig. 2A‘R’). The area surrounded by the cell membrane (CM) and ectocyst (EC) was measured using ZEN 3.3 blue edition software (Carl Zeiss Co. Ltd.) and an area ratio [area surrounded by CM (μm^2^) / area surrounded by EC (μm^2^)] was calculated.

For lectin staining, cysts over 2 weeks old were induced to excyst for 6 h. After excystment, the vacant cysts (i.e., ectocysts) were collected, washed three times with 1mM Tris-HCl and stained with an equal volume of 1 mg/mL wheat germ agglutinin (WGA): FITC conjugated (J-Oil Mills, Inc. Tokyo, Japan) at 4°C for 30 minutes. The lectin-stained ectocysts were washed three times with 1mM Tris-HCl and fluorescent staining was analyzed using an Axioscope A1 system with a 475 nm LED laser.

### Attenuated total reflectance infrared spectroscopy (ATR-IR) analysis

Two-week-old cysts were induced to excyst for 6 h and the ectocysts were collected, washed 3 times in distilled water and air-dried. The ATR-IR spectra of the ectocysts were recorded on a JOEL FTIR 6800 spectrometer equipped with a JOEL Pro One Diamond ATR unit. Each ectocyst was placed on the diamond prism and the spectra were recorded under a nitrogen atmosphere before and after washing with ethanol for 30 min. Powdered α-chitin was also measured on the prism to obtain a spectrum for chitin. A reference spectrum was taken with the prism without on any samples, and was subtracted from each sample spectrum during data processing.

### Bioassays

For the cell proliferation assay, *C. cucullus* vegetative cells were suspended at a low cell density (500 cells mL^-1^) in 4 mL of fresh Ex-medium containing different concentrations of NaCl (0, 0.3, 1.0, 3.5, 5.0, 10.0, 30.0% w/v). Cell numbers were counted at intervals after initiation of the culture; three aliquots of 20 μL were removed from the samples, cell numbers were counted under a Stemi 305 microscope (Carl Zeiss Co., Ltd., Tokyo, Japan), and calculated the density of cells was calculated.

Cell proliferative capability of excysted cells from cysts exposed to high salinity was evaluated by suspending the cells at a high density in En-medium and incubating them for 1 week. The suspension was then replaced with either En-medium containing 3.5% NaCl or fresh En-medium. After incubation for a week, the suspensions were replaced with Ex-medium and the cells were incubated for 6 h. Excysted cells were re-suspended in fresh Ex-medium at a low cell density (500 cells mL^-1^). Cell numbers were counted at hourly intervals in a 20 μL aliquot using a Stemi 305 microscope.

To evaluate the tolerance of cysts to salinity, cysts that were more than 1 week old were prepared in a Petri dish. The medium was replaced with En-medium containing different concentrations of NaCl (0, 0.3, 1.0, 3.5, 5.0, 10.0, 30.0% w/v), and the cyst samples were incubated as above. The exposed cysts were washed and re-suspended in a fresh Ex-medium for more than 6 h and an excystment assay was performed as previously described (27). To evaluate the effect of salinity on excystation, the rate of excystment of 1-week-old cysts, which were induced by Ex-medium with/without NaCl, was calculated.

The effect of saline exposure on encystation was measured as encystment (%) of exposed vegetative cells. The vegetative cells were induced by En-medium with or without 0.3% NaCl. The encystment rate was calculated as described previously (37).

Statistical analyses were performed using Tukey’s test in the Bell Curve for the Excel software package (Social Survey Research Information Co., Ltd., Japan).

### Total RNA extraction, library preparation, and de novo sequencing

Total RNAs were extracted from *C. cucullus* vegetative cells (300,000 cells) and 2-week-old mature cysts (3,000,000 cells) using the Direct-zol RNA purification system (Zymo Research Crop., California, USA) according to the manufacturer’s instructions. Paired-end cDNA libraries were constructed from 4 μg of total RNA using a KAPA Stranded mRNA-Seq Kit (Kapa Biosystems, Inc., Wilmington, USA) according to the manufacturer’s instructions. Samples included biological triplicates of cDNA. The integrity of the libraries was evaluated using a Bioanalyzer 2100 (Agilent Technology Japan, Ltd., Tokyo, Japan). The *de novo* sequencing (76 cycles, paired-end) was performed using Miseq (Illumina, K.K, Tokyo, Japan) at the Center for Bioscience Research and Education, Utsunomiya University.

### De novo transcriptome assembly and functional annotation

The raw sequence reads (76-bp read length) were processed using Trimmomatic (39,40) by trimming adapter sequences and low-quality ends (quality score, <15), and discarding the reads shorter than 50 bp. The obtained high-quality reads were further assembled into contigs (unigenes) using Trinity software (41). The sequences of unigenes were used for BLASTX searches and annotation against the Swiss-Prot protein database to predict biological functions (E-value cut-off was set at 1e-5). KEGG pathway analysis and Gene Ontology (GO) analysis for unigenes were performed using the KEGG automatic annotation server (KAAS) and InterproScan, respectively (42,43).

### Differentially expressed gene analysis

Differentially expressed genes (DEGs) were identified as described previously (44). rRNA sequences were excluded from Trinity unigenes by removing matching entries in the rRNA database using the megablast program. High-quality reads derived from each sample were further mapped to rRNA to remove transcripts. Gene expression was calculated using the Fragments Per Million Reads (FPKM) method, and *P*-values were determined by the false discovery rate. DEGs were identified as 2-fold up- or down-regulated with a false discovery rate (FDR) < 0.05 between vegetative cells and cysts in this study.

### Real time PCR analysis

Total RNAs were isolated from vegetative cells, 2-week-old cysts, and encystment-induced cells as described above. RNA was reverse transcribed using a Transcriptor First Strand cDNA synthesis Kit (Roche Diagnostics, Mannheim, Germany) according to the manufacturer’s instructions. Gene-specific primers were designed using Primer3 software (table S1). All reactions were performed using the Real time PCR system STEP1 (Thermo Fisher Scientific K.K., Tokyo, Japan) with Power up SYBER (Thermo Fisher Scientific) and three technical replicates. For verification of the RNA-Seq, amplification was performed using the following protocol: 95°C for 10 min, followed by 40 cycles at 95°C for 15 sec and 60°C for 1 min. Melt curves were produced according to the manufacturer’s instructions. The amplification conditions for analysis of the *CHS5* gene were as described in the manufacturer’s instructions. All data were analyzed by the ΔΔct method using Real-Time PCR System Software (Thermo Fisher Scientific). Transcription elongation factor *SPT4* or the α-tubulin gene of *C. cucullus* (45) were selected as internal control genes. Primers were designed using Primer 3 Plus software (https://www.bioinformatics.nl/cgi-bin/primer3plus/primer3plus.cgi) for the regions shown in table S3.

### Endocyst protein assay

Excystment was induced overnight in cyst samples (10^6^ cells) using 1% rice leaf infusion without protein (CMP-; proteins including in the infusion were removed using a 3-kDa protein cut filter). The empty ectocysts adhering to petri dishes were discarded and the supernatants containing medium soluble endocysts were collected. Before the collection, the excysted cells (vegetative cells) were removed with a 1 μm pore size membrane filter to prevent contamination of proteins of excysted cells, but 3×10^3^ cells/100mL remained in the sample. The endocyst proteins precipitated by 4 times the volume of acetone at −20°C overnight, centrifuged at 15,000 g at 4°C for 15 min, air-dried, and the pellet was re-suspended in Tris-HCl buffer. The samples were concentrated using a 3-kDa cut filter and solubilized with 4-strength SDS-Page sample buffer. To determine protein contamination by CMP- and contaminated excysted cells, the same volume of CMP- and excysted cells in the endocyst samples, were processed in the same way. Additionally, vacant ectocyst samples were prepared to determine protein contamination by ectocysts. The vacant ectocysts corresponding to 10^6^ cells were collected, disrupted using a beads homogenizer and ultrasonic homogenizer, and centrifuged (15,000 x g at 15 min). The pellet was solubilized in SDS-sample buffer and boiled for 3 min.

SDS-PAGE and western blotting on PVDF membranes were performed as described previously (37). Enzymatic digestion of proteins on PVDF membranes was as previously described (46). Briefly, bands were excised from the membrane, rehydrated in 100 µl ultrapure water, and the pieces of membrane were cut into 1 mm-wide squares. The membrane pieces were placed in a digestion buffer consisting of 1% hydrogenated Triton X-100, 10% acetonitrile, and 100 mM Tris-HCl (pH 8.0) and incubated for 30 min at room temperature. Then, a 1:10 ratio of 0.1 µg/µl of trypsin (Thermo Fisher Scientific) was added to the digestion buffer and the samples were incubated for 24 h at 37°C. After the incubation period, the samples were sonicated twice for 5 min and 50 µl of digestion buffer was added. The supernatant was transferred to a new tube, and 100 µl 0.1% trifluoroacetic acid was added. Finally, 2 µl 1% diisopropylfluorophosphate was added. The extracted peptides were cleaned using Spin Tip columns (C18 resin; Thermo Fisher Scientific) according to the manufacturer’s desalted protocol, with a final elution of 20 µl 2% acetonitrile and 0.1% formic acid. The concentration of purified peptides was measured using a Nanodrop Spectrophotometer (Thermo Fisher Scientific). After protein extraction and purification, the protein concentrations of the samples ranged from 0.98-1.4 mg/ml. Peptides produced by protease digestion were separated with a Thermo UltiMate 3000 UHPLC (Thermo Fisher Scientific) using a 5–80% effective gradient with mobile phase B (98% acetone, 0.1% formic acid) for 60 minutes. The separated peptides were ionized using a nanoESI source and then subjected to tandem mass spectrometry in a Q-Exactive HF X (Thermo Fisher Scientific). The LC-MS/MS analysis and gene ontology analysis were performed at the Beijing Genomics Institute, China. The results were processed by PEAKS online ver. 10 and obtained peptide sequences were searched in our RNA-seq data base (DRR275117-DRR275118.) in which RNA sequences have been translated into amino acid sequences.

## Results

### Bioassay

*C. cucullus* resting cysts were able to survive in 3.5% NaCl, a salt concentration comparable to seawater, for a week and the cells maintained their proliferative ability. However, vegetative cells died immediately on suspension in high salinity media (1.0–30%) (Fig. 1A). Vegetative cells survived more than 24 h in 0.3% saline (Fig. 1A) but could not form cysts (Fig. 1B) or proliferate; they gradually died after more than 24 h in this medium (Fig. 1A). One-week-old cysts survived in 10% salinity for 12 h (Fig. 1C) and showed an excystment rate of 80.9%. A low rate of encystment (0.6%) was found when they were exposed to 30% salinity for 12 h. *Colpoda* resting cysts survived in 3.5% salinity for more than 1 week (Fig. 1D) and showed an excystment rate of 80.1%. After exposure, the resting cysts were able to excyst and proliferate similarly to the control group although at a slower rate (Fig. 1E). Resting cysts treated with 0.3% salinity could excyst and revert to the vegetative state under this saline condition, whereas those suspended in 0.5–30% saline did not show excystment (Fig. 1F).

**Fig. 1.**
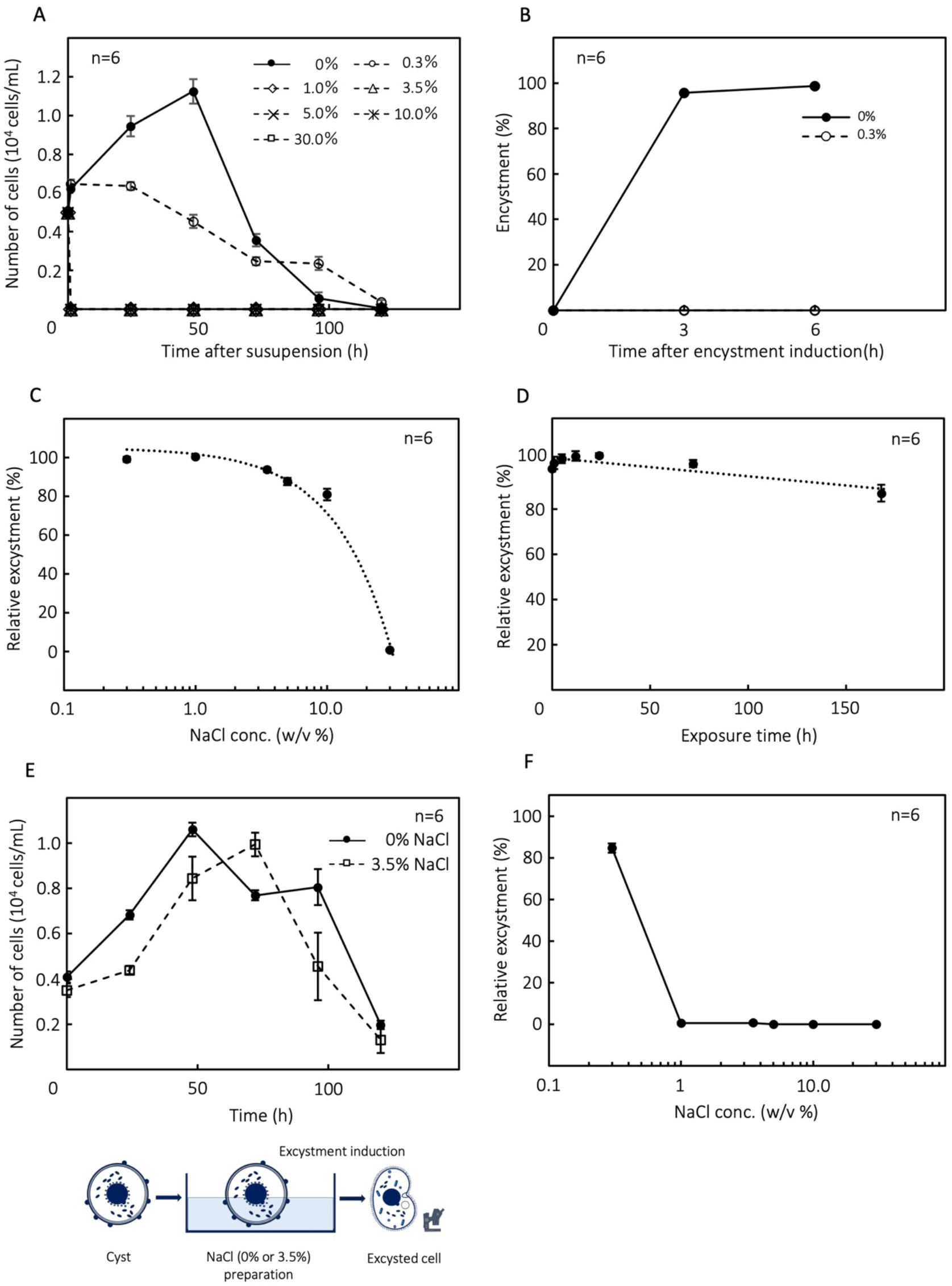
Bioassay of *Colpoda* tolerance to different salinity conditions (0, 0.3, 1, 3.5, 5, 10, 30% NaCl medium). (A) Vegetative cell tolerance to saline medium. (B) Encystment assay in 0 and 0.3% saline medium. (C) Tolerance of *Colpoda* cysts to exposure to saline media for 12 h. (D) Tolerance of *Colpoda* cysts to 3.5% saline medium for 0–168 h. (E) Cell proliferation assay of excysted cells from cysts exposed to 0 or 3.5% NaCl medium. (F) Rate of excystment assay of cysts in different saline media (0, 0.3, 1, 3.5, 5, 10, 30% NaCl medium).

**Fig. 2.**
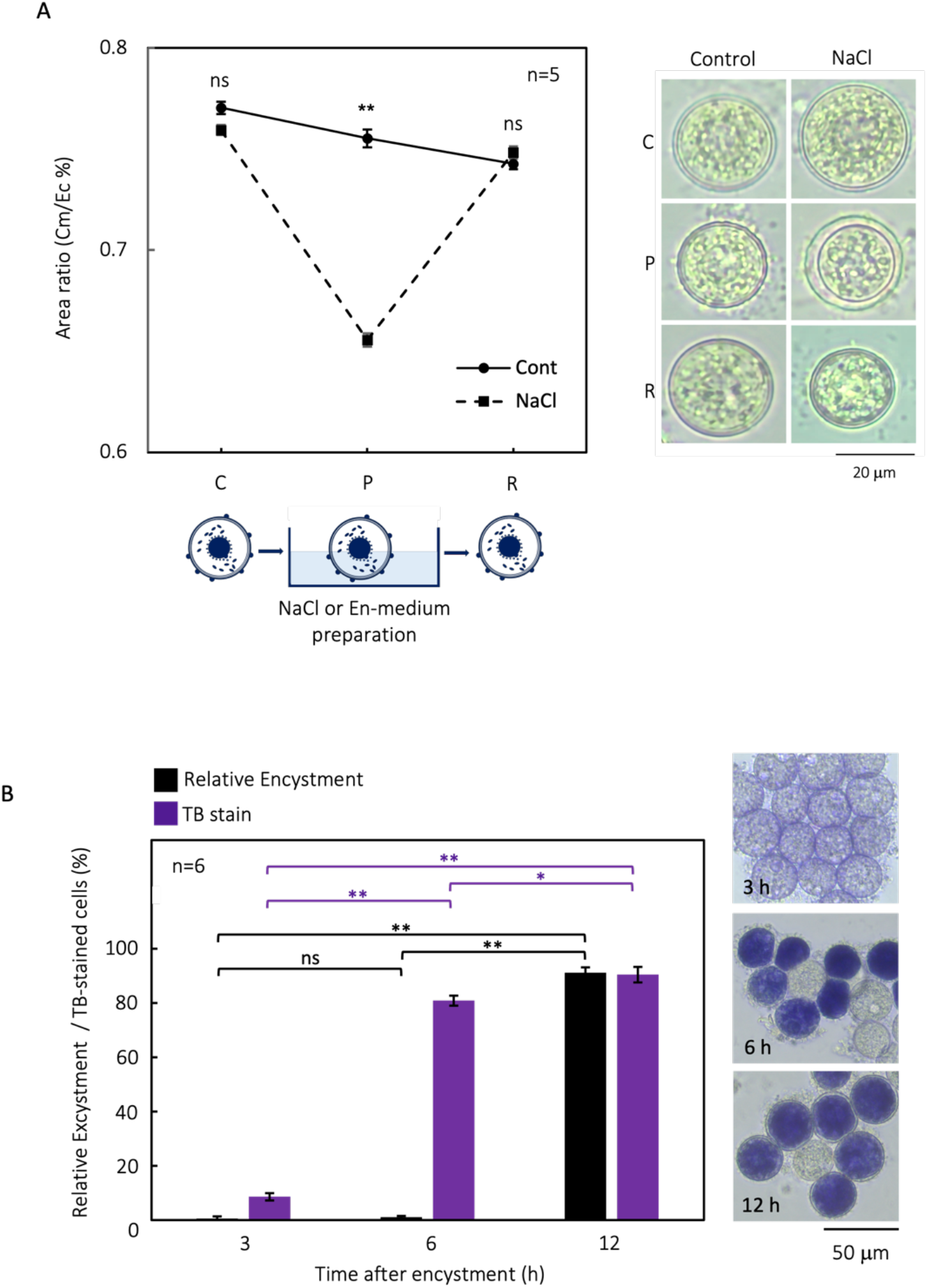
Influence of salinity on *Colpoda cucullus* morphology. (A) Effect of saline medium on the cell membrane. The area ratio and micrographs of 1-week-old cysts suspended in 3.5% NaCl (C) or control El medium for 1 week (P), and resuspended in El medium (R). (B)Micrographs show TB staining at 3, 6, and 12 h after the onset of encystment induction. The rate of TB staining and relative excystment rate at 3, 6, and 12 h after induction of encystment in cysts that had been suspended in 3.5% NaCl for 1 h. * p < 0.05; ** p < 0.01; ns, no significant difference.

### Morphological analysis

Our morphological analysis indicated that salinity tolerance in cysts was due to the formation of the cell membrane. Vegetative cells were impaired immediately after being suspended in a high-salinity medium, but cysts remained viable. When 1-week-old cysts (Fig. 2A ‘C’) were suspended in 3.5% saline for 1 week (Fig. 2A ‘P’), there was clear shrinkage inside the cyst; however, this recovered when the cysts were returned to the normal medium (Fig. 2A ‘R’). Hence, we conclude that the cell membrane was not affected by high salinity and protected the cell from its adverse effects.

TB staining showed that the endocyst was not involved in the high salinity tolerance of resting cysts. At 3 h after the induction of encystment, <10% of the cells were TB-stained; by contrast, over 80% of cells were TB-stained at 6 h and 12 h (Fig. 2B). The relative excystment rates in high salinity (3.5% NaCl for 1 h) were not significant at 3 h and 6 h (≤1%, respectively) but were significantly elevated at 12 h (>90%; Fig. 2B). Overall, there was no correlation between salinity tolerance and the rate of endocyst formation.

### Transcriptomic profiles of vegetative cells and cysts

Paired-end sequences (2 × 75 bp in length) from *C. cucullus* vegetative cells and cysts were generated by Miseq, which produced 13,518,200 reads and 12,733,840 reads, respectively (Table SM1). The de novo assembly results are summarized in table S2. In total, we obtained 73,011 (N50 length = 1,443 bp) unigenes in *C. cucullus* (table S2). The total length was 65,639,138 bp, and the maximum sequence length was 17,589 bp; the minimum sequence length was 201 bp (table S2). We found that the highest number of unigenes was represented in groups 201–300, and the lowest in group 1901– 2000 (fig. S1). All of the transcriptome data are provided in table S4 and the sequence reads have been deposited in the DDBJ Sequence Read Archive (DRA)/SRA under the accession numbers DRR275117-DRR275118.

### MA and volcano plot analysis

We used an MA plot to compare the log_2_ fold-changes (Log_2_ FC) of unigenes in vegetative cells and cysts and mean expression (log counts) (Fig. 3A). Overall, 14,056 differentially expressed unigenes (DEGs) were identified in vegetative cells (FDR <0.05, Log_2_FC >1) and 11,258 in cysts (FDR <0.05, −Log_2_FC >1); these are shown as red points in the MA plot (Fig. 3A). A comparison of P-value and log fold-change between vegetative cells and cysts is shown in the volcano plot (Fig. 3B). DEGs in the vegetative cells and cysts (FDR <0.05, Log_2_FC >1) are shown as red points.

**Fig. 3.**
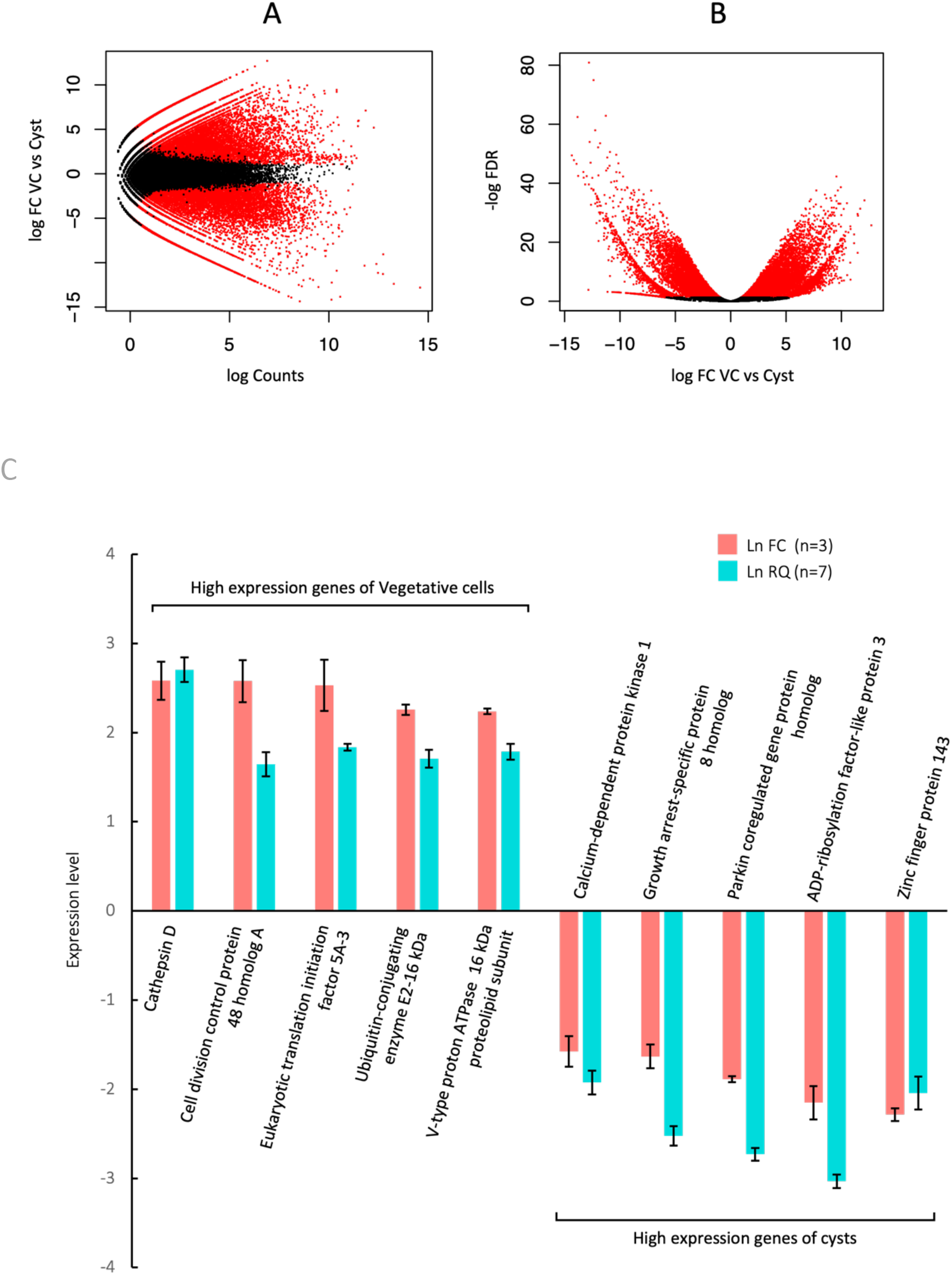
MA plot, Volcano plot, and verification of the RNA-Seq data by qRT-PCR analysis in *C. cucullus* vegetative cells and cysts. (A) MA plot of DEGs. The x-axis represents the log2 transformed mean expression level of unigenes. The y-axis represents the log2 transformed fold change in vegetative cells vs cysts. The positive or negative value indicates vegetative cell-specific or cyst-specific, respectively. (B) Volcano plot of DEGs. The x-axis represents the log2 transformed fold change in vegetative cells vs cysts. The positive or negative value indicates vegetative cell-specific or cyst-specific, respectively. The y-axis represents the log2 transformed fold change. P < 0.05, FC >1 genes are indicated by red points. (C) Verification of the RNA-Seq (Ln FC) data by qRT-PCR analysis (relative gene expression quantity; Ln RQ). Five genes with high expression in vegetative cells and cysts were selected. FC (n = 3) corresponds to the expression level of the transcriptomic analysis and RQ (n = 7) is that of qRT-PCR.

### Verification of RNA-Seq data

The expression levels of five up-regulated genes in vegetative cells and cysts were verified by real-time PCR. The mRNA expression levels of these genes were consistent with the transcriptomic data (Fig. 3C). Transcription elongation factor *SPT4* was used as the internal control as its expression is similar in vegetative cells and cysts. The names of the selected genes are given in Fig. 3C and their gene IDs and amplification primers are shown in table S3.

### Functional annotation of DEGs and GO enrichment analysis

All unigenes were searched against the Swiss-Prot database; the identifies of 23,265 unigenes were obtained. Unigenes that were over 2-fold up- or downregulated were isolated: 14,056 genes were expressed significantly higher in vegetative cells, whereas 11,258 genes were expressed significantly higher in resting cysts (table S4).

We performed GO enrichment analysis of vegetative cells and cysts and characterized the 98,560 unigenes as 897 GO terms. These GO terms could be divided into 374 categories of Biological Process, 383 categories of Molecular Function, and 140 categories of Cell Components (table S5). Expression levels were significantly different between vegetative cells and cysts for 28 Biological Process categories with 6,691 unigenes, 32 Molecular Function categories with 5,215 unigenes, and 9 Cell Components categories with 498 unigenes. In vegetative cells, unigenes of GO category ‘cysteine-type peptidase activity’ showed the greatest upregulation, followed by ‘chromatin binding’, and ‘hydrolase activity, acting on acid anhydrides, in phosphorus-containing anhydrides’ (Fig. 4). In cysts, unigenes of the GO category ‘phosphorelay sensor kinase activity’ showed greatest upregulation, followed by ‘phosphorelay signal transduction system’ and ‘signal transduction’ (Fig. 4). In vegetative cells (Fig. 5A), the number of unigenes in the GO category ‘integral component of membrane’ was the largest and similarly for the GO Cell Components type. The GO ‘nucleus’ and ‘DNA binding’ categories also contained many of the unigenes: the number of unigenes in ‘DNA binding’ and ‘proteolysis’ were the largest in Molecular Function and Biological Process, respectively. In cysts (Fig. 5B), the GO ‘membrane’ category was the largest and all of them are included in the categories GO category Cell Components. The GO ‘signal transduction’ and ‘phosphorelay signal transduction system’ categories also included many unigenes: ‘signal transduction’ and ‘phosphorelay sensor kinase activity’ were the largest categories in Biological Process and Molecular Function, respectively. Interestingly, 47% of unigenes identified in cysts fell into the ‘membrane’ category.

**Fig. 4.**
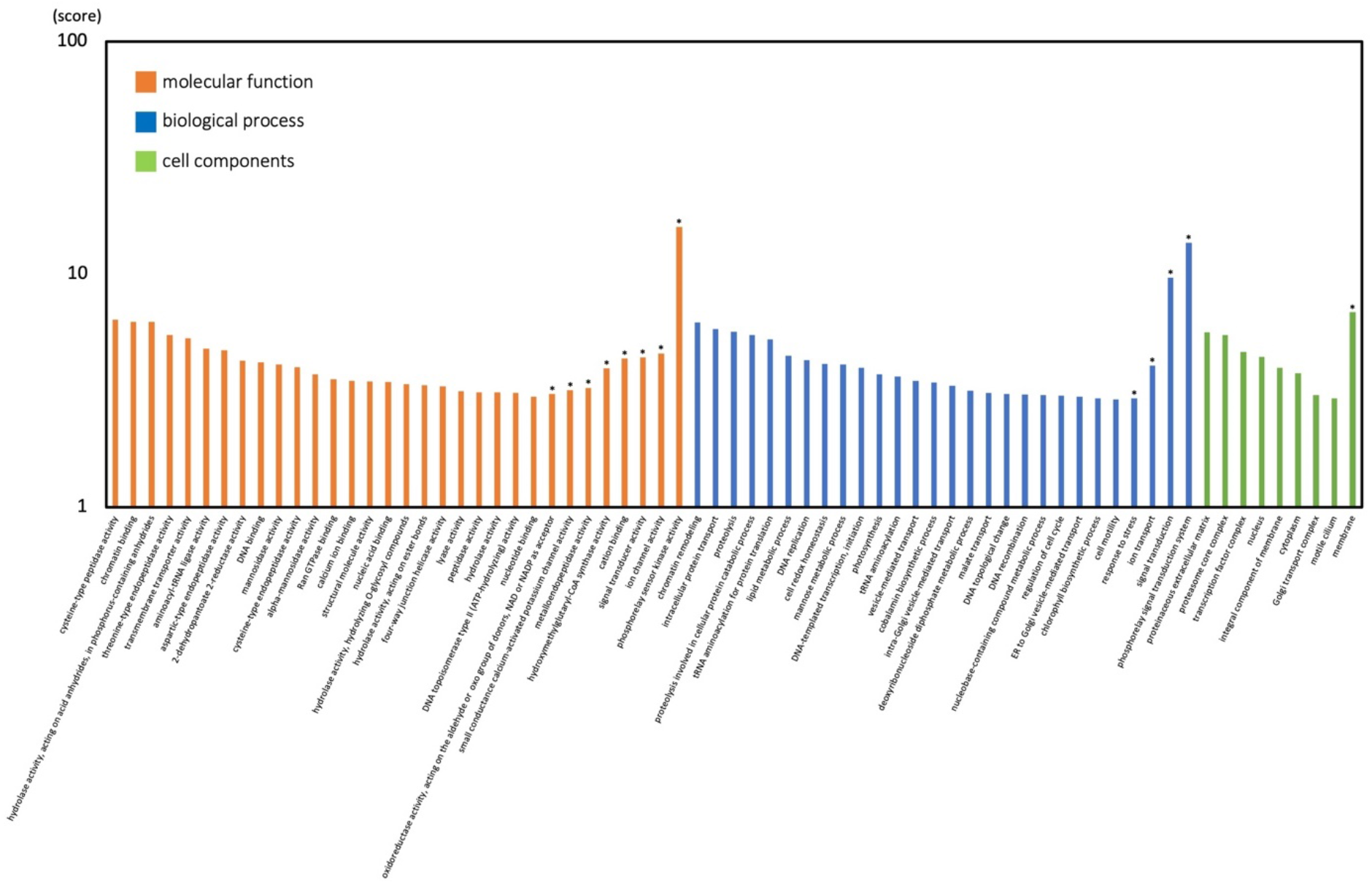
GO functional classification of differentially expressed unigenes. Z score of GO terms is shown as the y-axis. Asterisks indicate cysts.

**Fig. 5.**
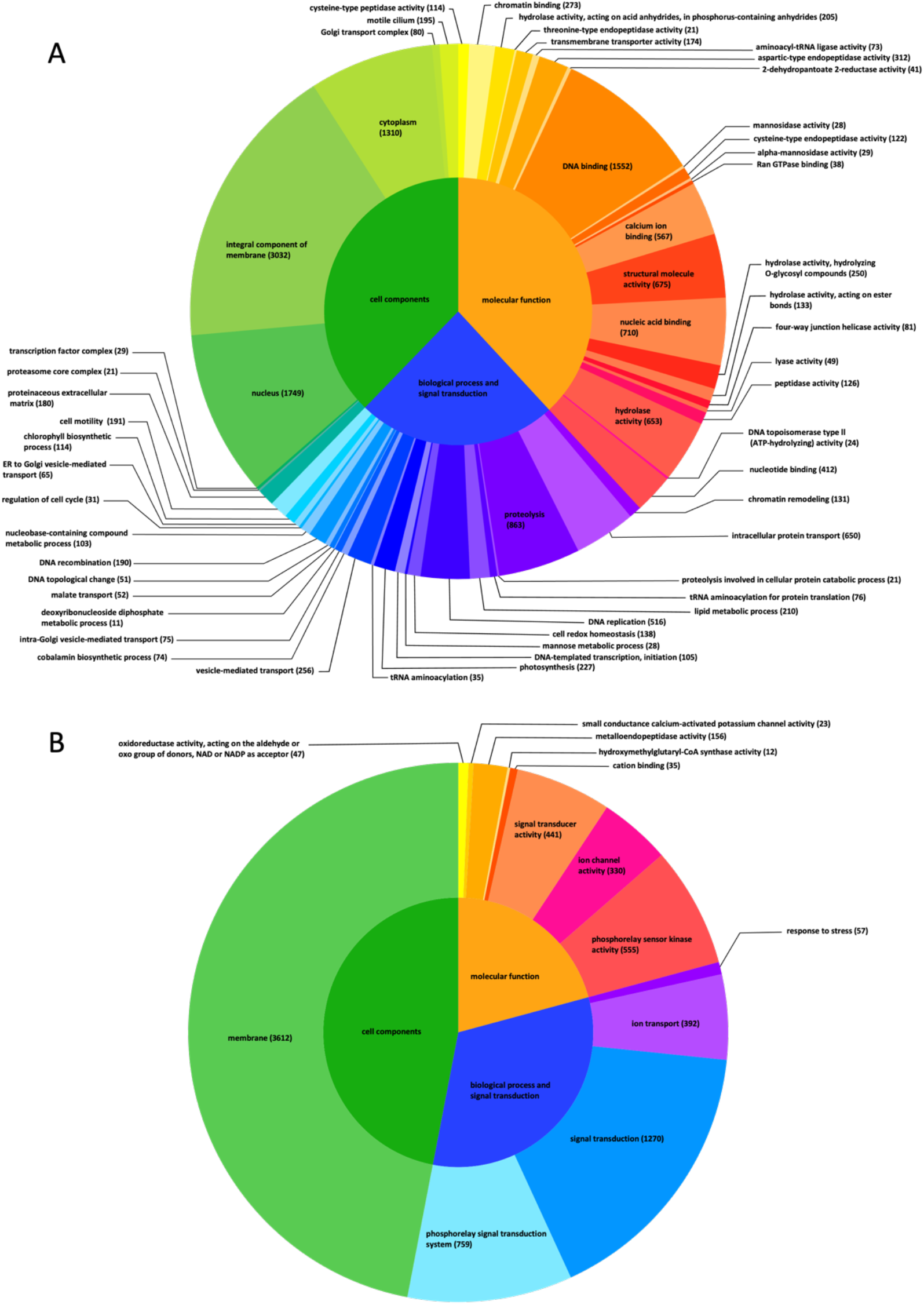
Gene ontology enrichment analysis of the DEGs in vegetative cells (A) and cysts (B)

### Analysis of the cyst wall: ectocyst components and endocyst proteins

An excysting cell and the two main layers of the cyst wall are shown in Fig. 6A. As our analyses have indicated that cyst wall components are likely to be involved in salinity tolerance, we sought to identify them. Previous studies have revealed many of the protein components of the cyst wall (ectocyst) (47–49) and the presence of elongation factor Tu (EF-Tu) in lepidosomes (50) in the genus *Colpoda*. However, there is no information to date on purified endocysts. Here, endocyst proteins (En) were purified and electrophoresed by SDS-PAGE (Fig. 6B); the figure also shows proteins from protein-free culture medium (CMP-), vegetative cells, and ectocysts (EC). SDS-PAGE indicated that the endocyst sample was unaffected by CMP- and vegetative cells. The endocyst-specific protein bands (p48, p36, p21a, p21b, p19, and p17), which were not present in ectocysts, were analyzed by LC-MS/MS and a summary of the results is shown in Table 1. Bands p48, p36, and p19 were predicted to be proteins homologous to actin; p48 also contained elongation factor 1 alpha. In addition, some enzymes, membrane proteins and transmembrane proteins were detected.

**Fig. 6.**
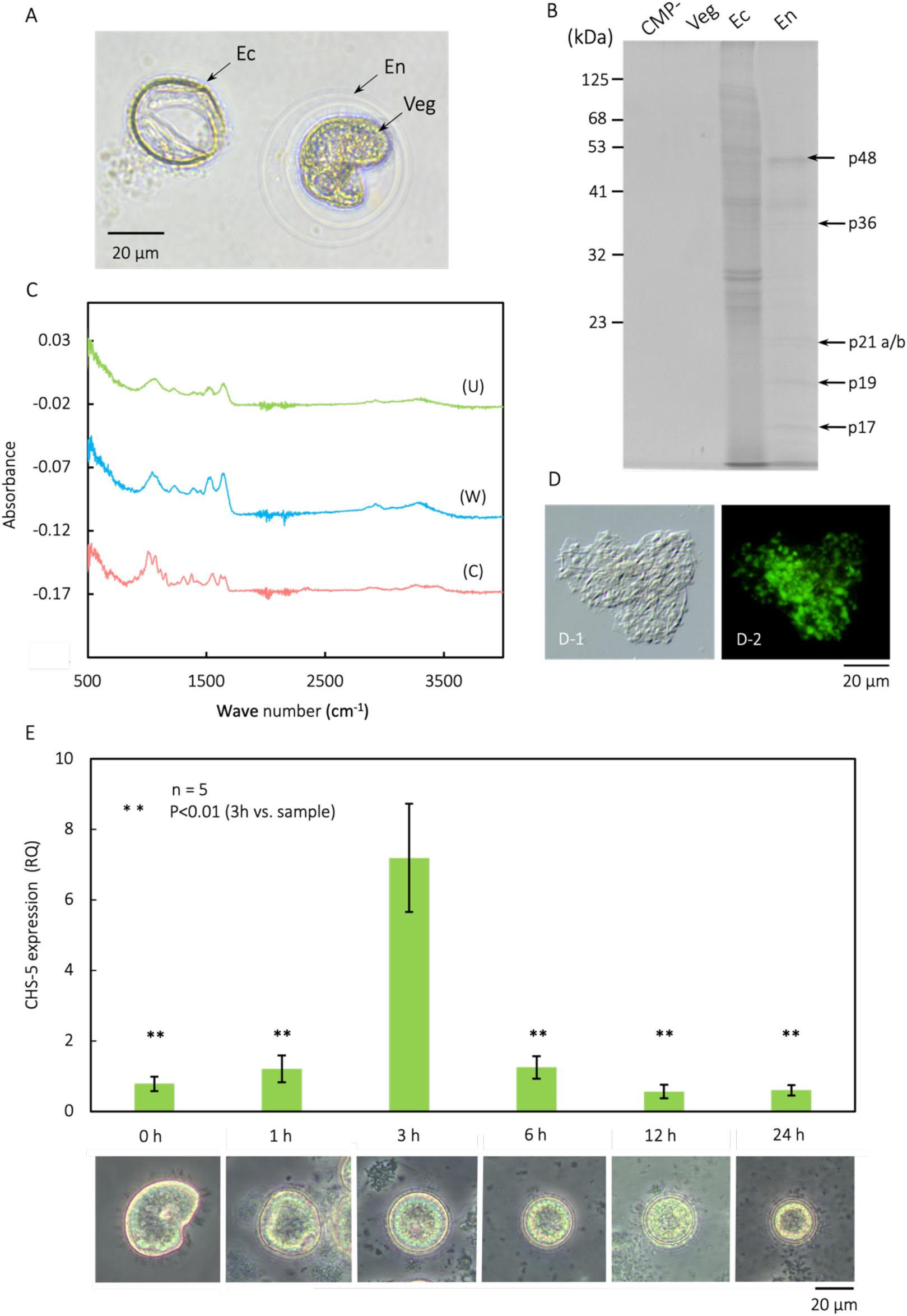
Analysis of cyst wall (endocyst and ectocyst) components. (A) Light microscope images of an excysting vegetative cell (Veg) and the main two layers of cyst wall: endocyst (En) and ectocysts (Ec). (B) Identification of endocyst-specific proteins. SDS-PAGE of proteins from endocysts (En), ectocysts (Ec), and vegetative cells (which seem to show some contamination by the endocyst sample (Veg), and culture medium without protein (CMP-). Endocyst-specific protein bands (p48, 36, 21a/b, 19, and 17) were identified by LC-MS/MS and the results are shown in Table 1. C-ATR-IR spectra of an unwashed ectocyst sample (U), an ethanol-washed ectocyst sample (W), and pure chitin sample (C). An overview of absorbance peaks is shown in Table 2. (D) Lectin staining (FITC-conjugated WGA) of an ectocyst sample by Differential Interference Contrast (DIC) microscopy (D-1) and fluorescence microscopy (D-2). The bar represents 20 μm. (E) Morphological changes in the encystation process of *C. cucullus* and qRT-PCR analysis of expression of *CHS5* gene during this process. ** p < 0.01 versus the 3 h sample.

**Table 1.**
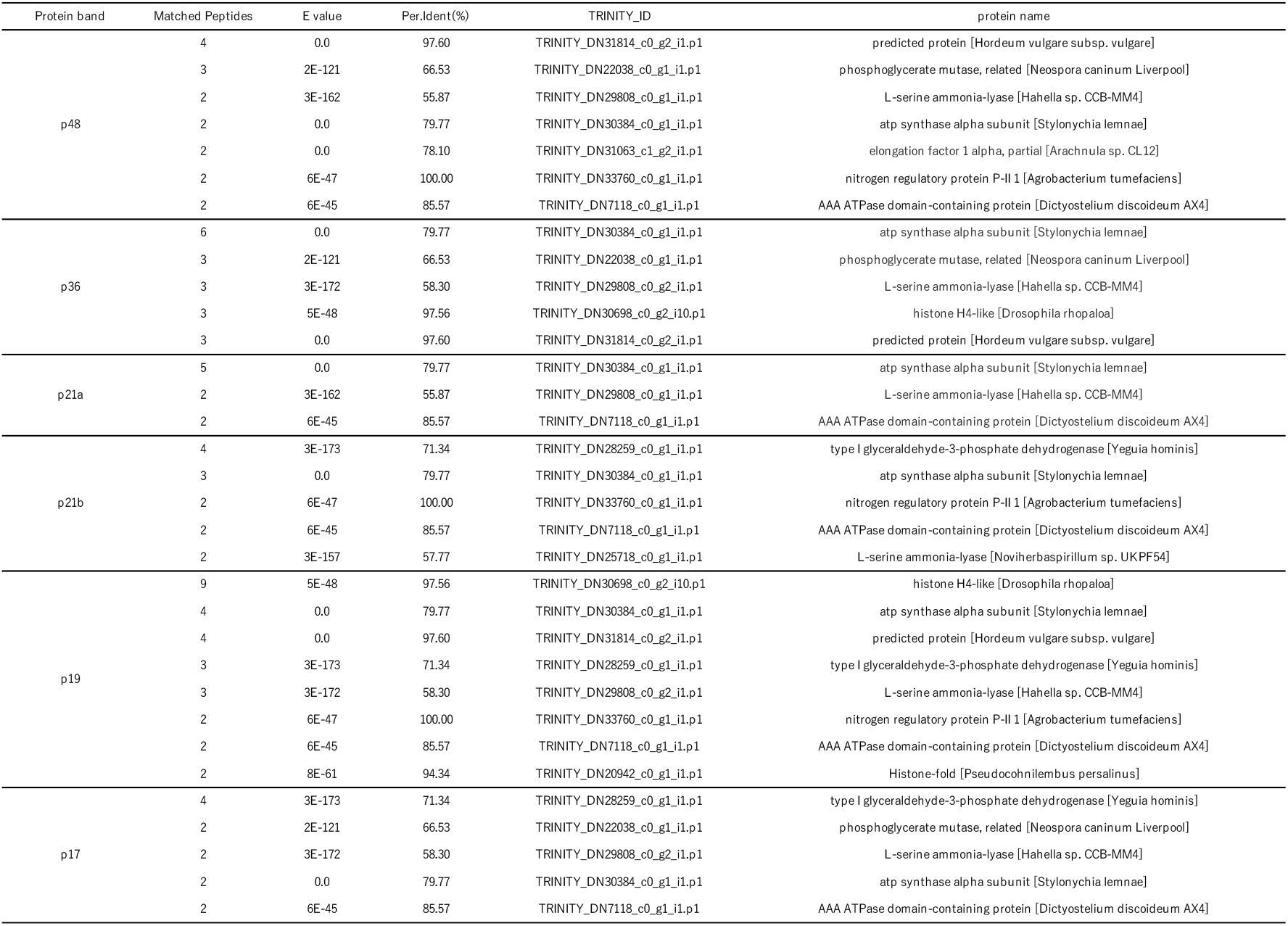
Identification of endocyst specific protein bands (p48, 36, 21a/b, 19, and 17) by LC-MS/MS analysis.

**Table 2.**
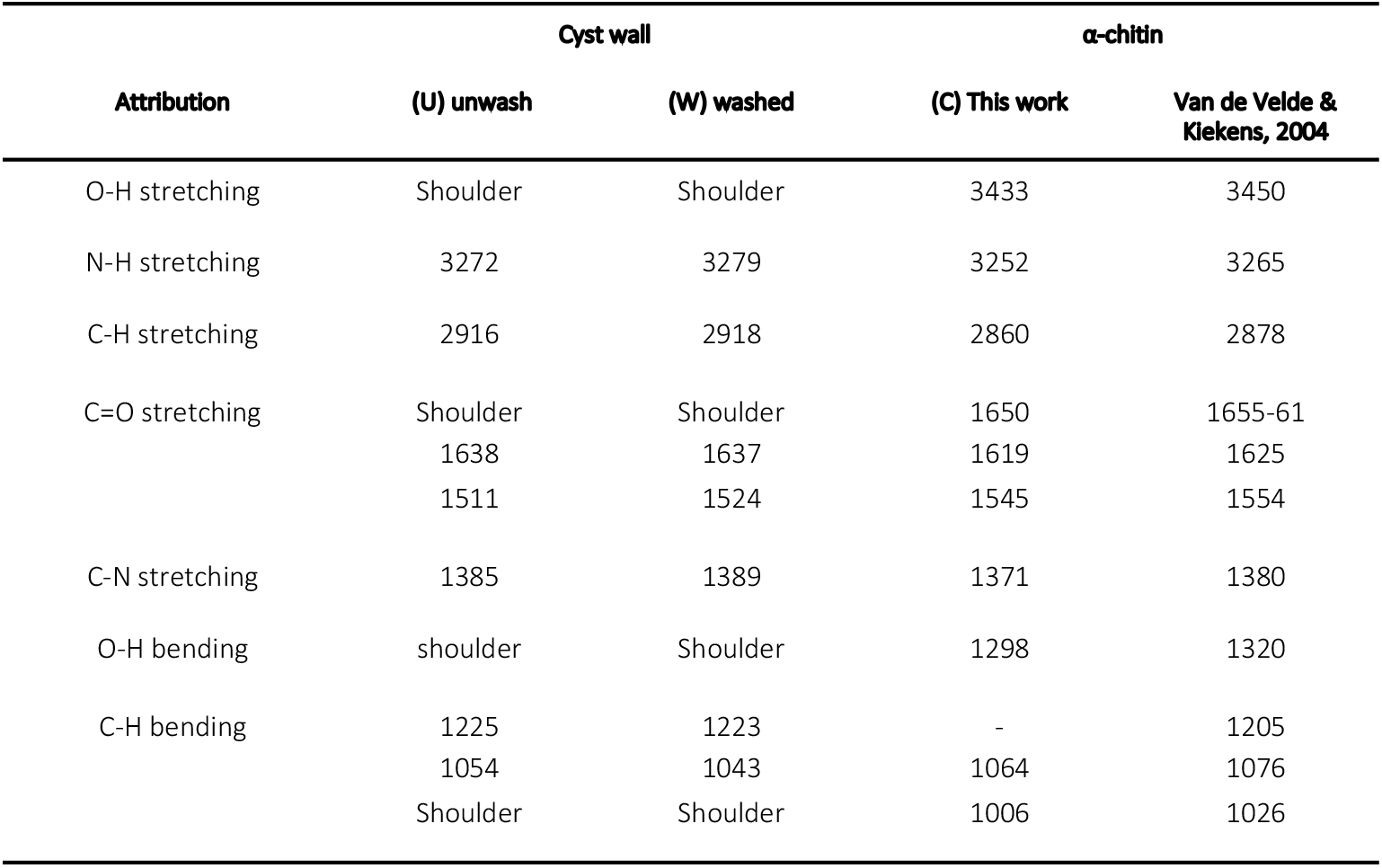
Overview of absorbance peaks of ATR-IR spectra of cyst wall and α-chitin samples. The spectra peaks of unwashed ectocyst sample ’U’, ethanol washed ectocyst sample ’W’, pure chitin sample ’C’and that by Van de Velde and Klekens, 2004 (51) are shown.

| Attribution | Cyst wall | | α-chitin | |
| --- | --- | --- | --- | --- |
|  | (U) unwash | (W) washed | (C) This work | Van de Velde & Kiekens, 2004 |
| O-H stretching | Shoulder | Shoulder | 3433 | 3450 |
| N-H stretching | 3272 | 3279 | 3252 | 3265 |
| C-H stretching | 2916 | 2918 | 2860 | 2878 |
| C=O stretching | Shoulder | Shoulder | 1650 | 1655-61 |
|  | 1638 | 1637 | 1619 | 1625 |
|  | 1511 | 1524 | 1545 | 1554 |
| C-N stretching | 1385 | 1389 | 1371 | 1380 |
| O-H bending | shoulder | Shoulder | 1298 | 1320 |
| C-H bending | 1225 | 1223 | - | 1205 |
|  | 1054 | 1043 | 1064 | 1076 |
|  | Shoulder | Shoulder | 1006 | 1026 |

The spectra obtained in an ATR-IR analysis indicated that the cyst wall flake contained chitin. For ectocysts, the spectra from the ATR-IR analysis were similar to that of α-chitin (Fig. 6C) (51) except for the O-H stretching region (3100-3600 cm^-1^). The two broad OH peaks of α-chitin (3433 cm^-1^, 3252 cm^-1^; Fig. 6C) were present as a single broad peak (3272 or 3279 cm^-1^) in the unwashed and ethanol-washed ectocyst samples (Fig. 6C); the difference between them may be due to interaction between chitin and other components in the wall as the OH stretching modes might be significantly influenced by hydrogen bonding with other components such as oil and proteins. There was an additional small difference between the chitin sample and the ectocyst samples under 3100 cm^-1^ and the difference can again be explained by the effect of other components such as acetylation or crystallinity. The small differences between the ethanol-washed and unwashed samples demonstrate that some other components were removed by ethanol. The main peaks and their attribution are summarized in Table 2.

The results of the lectin (WGA) staining supported those of the ATR-IR analysis. Ectocyst samples (Fig. 6D-1) were stained with FITC-labeled WGA, a lectin that binds with chitin (52); an FITC signal was detected in the ectocysts (Fig. 6D-2), indicating the presence of chitin in the ectocyst.

*CHS5* is involved in chitin synthesis in *Saccharomyces cerevisiae* (53) and has been shown to have a temporary increase in expression during ectocyst synthesis (Fig. 6E). The ectocyst is gradually formed during encystation and can be observed at 3 h after encystment induction (Fig. 6E ‘3 h’). We performed real-time PCR analysis and showed that expression of the *CHS5* gene was increased temporarily at 3 h after the onset of encystment in *Colpoda*.

## DISCUSSION

The resting cysts of *C. cucullus* demonstrate pronounced salinity tolerance that might allow the protist to be dispersed by oceanic currents. The salinity tolerance of this soil ciliate is higher than that of soil testate amoebae; the adaptation of testate amoebae to rising sea levels was demonstrated in a study from Tuvalu (54,55).

Salinity tolerance in *Colpoda* is acquired by the extensive membrane reconstruction associated with encystation, as revealed here by morphological and gene ontology analyses. The properties of the membrane change considerably and these changes probably enable it to maintain its structure in high salinity conditions. The gene annotation analysis indicated that unigenes belonging to Molecular Function and Biological Process were downregulated, whereas those in the ‘membrane’ category of Cell Components were upregulated in resting cysts. DEGs of the ‘membrane’ (Cell Components) and ‘ion channel activity’ (Molecular Function) were abundant in cysts, thus reconstruction of the cell membrane was activated during encystment. In resting cysts, the Biological Process category contained a large proportion of the identified DEGs, whereas gene variety of was simplified. The expression of genes related to stress responses and oxidoreductase activity showed upregulation, while other metabolic processes displayed down-regulation. Our results of this study support our previous claim that resting cysts are not just resting but repairing cell damages even though they cease mitochondrial metabolic activity (56), i.e., carbonylated proteins induced by gamma irradiation have been shown to be repaired in resting cysts (37). Differential gene expression during the early stage of encystment has been shown previously in transcriptomic studies of *Colpoda aspera* (57) and *Pseudourostyla cristata* (58); these observations demonstrated the presence of an early signaling pathway for encystment as had been suggested earlier (59,60). Encystment in *C. cucullus* is controlled by a complex pattern of gene expression. Thus, 14,056 unigenes were differentially expressed in vegetative cells, and 11,258 unigenes were differentially expressed in mature cysts. The differential gene expressions lead to morphological changes, e.g., the characteristic broad bean cell shape gradually becomes spherical (61), a cyst wall is formed, cilia and mitochondria are digested (62).

One important component of the ectocyst is chitin. The synthesis of chitin in *Colpoda* as an ectocyst component was shown here by AT-IR, WGA (lectin) staining, and activation of *CHS5* expression. Carbohydrates such as cellulose in plants and chitin in invertebrates coat cell surfaces. The cyst walls of ciliates also contain carbohydrates (24), and the presence of chitin has been inferred in some ciliates following chitinase digestion (24, 63, 64). However, *Colpoda* species (*C. cucullus, C. steinii*) have been reported to test negative for chitinase digestion, indicating the absence of chitin (24,47,63). On the other hand, we confirmed the presence of chitin in ectocysts by chemical analysis in this study. In previous studies, it is possible that the enzymatic activity of chitinase was inhibited in some manner, possibly involving highly acidic proteins (24). Our method is completely unaffected by that. Although chitin has been found in the cyst walls of many ciliates, chitin synthase activity has not yet been reported (24). The *CHS5* gene is not the chitin synthase itself but is considered to be involved in chitin synthesis as knockout of *CHS5* expression inhibits chitin synthesis (53). Hence, the result that activation of *CHS5* expression coincided with formation of ectocysts supports the interpretation that chitin is a components of ectocysts.

In addition to the ectocyst acting as a chitinous biological armor, the inner endocyst is likely to serve as another protective barrier by shielding the cell from environmental stresses such as desiccation and dehydration. Actin was detected in protein bands of p48, 36, and 19 as possibly the main components of the endocyst, and EF-1 alpha was also detected in protein bands of p48 as possibly one of its components. Actin is likely to be in the form of F-actin as ectocyst and endocyst cell walls and lepidosomes can be stained with Phalloidin (65). However, it is possible that this staining may be partially attributable to autofluorescence of the endocyst layer (65). The presence of endocyst actin is unlikely to be a contaminant from ectocysts and/or lepidosomes, as actin has not been detected in samples of ectocysts. Most probably, the actin originated from endocysts. However, it is difficult to completely eliminate contamination of intracellular actin with the purification method used for endocysts in this study. A similar limitation may apply to other detected proteins for example, the histones and membrane proteins such as ATP synthase alpha subunit that were detected in some protein bands of endocysts. In contrast, it is possible that some proteins of the cell are included and that they are materials used to construct the endocyst since endocyst layers are formed by the exocytosis of short filamentous precursor materials (62). The fact that endocyst layers stained with TB (34, 62) indicates that they include mucopolysaccharides. The level of EF-1 alpha has been reported to be elevated during encystment, (66) and may contribute to endocyst formation by polymerization of actin (67). On the other hand, the level of EF-1 alpha is reduced within 1 h of excystment induction (68). This process is likely dependent on actin depolarization. The actin identified in this study had a molecular weight different from the expected 42 kDa. This is probably due to the influence of post-translational modification, such as acetylation and phosphorylation, as has been observed in *Entamoeba histolytica* (69), or fragmentation due to the use of trypsin. Various molecular weights of actin were detected from Hela cell protein samples (70). Other membrane proteins may also function as a second biological barrier by confining water or acting like a polymer absorber to maintain humidity and protect against desiccation.

The data presented here support our proposal of oceanic dispersal of freshwater protists as shown in Fig. 7A. It is possible that they could cross oceans as floating cysts, being protected by a cyst wall consisting of a chitinous outermost layer (ectocyst) and several inner layers (endocyst) of actin-like proteins with mucopolysaccharides. The cysts would be passively transferred by oceanic currents which enable long-distance transportation (71). This oceanic dispersal might be complemented by other means such as transport by insects, birds, or human activities.

**Fig. 7.**
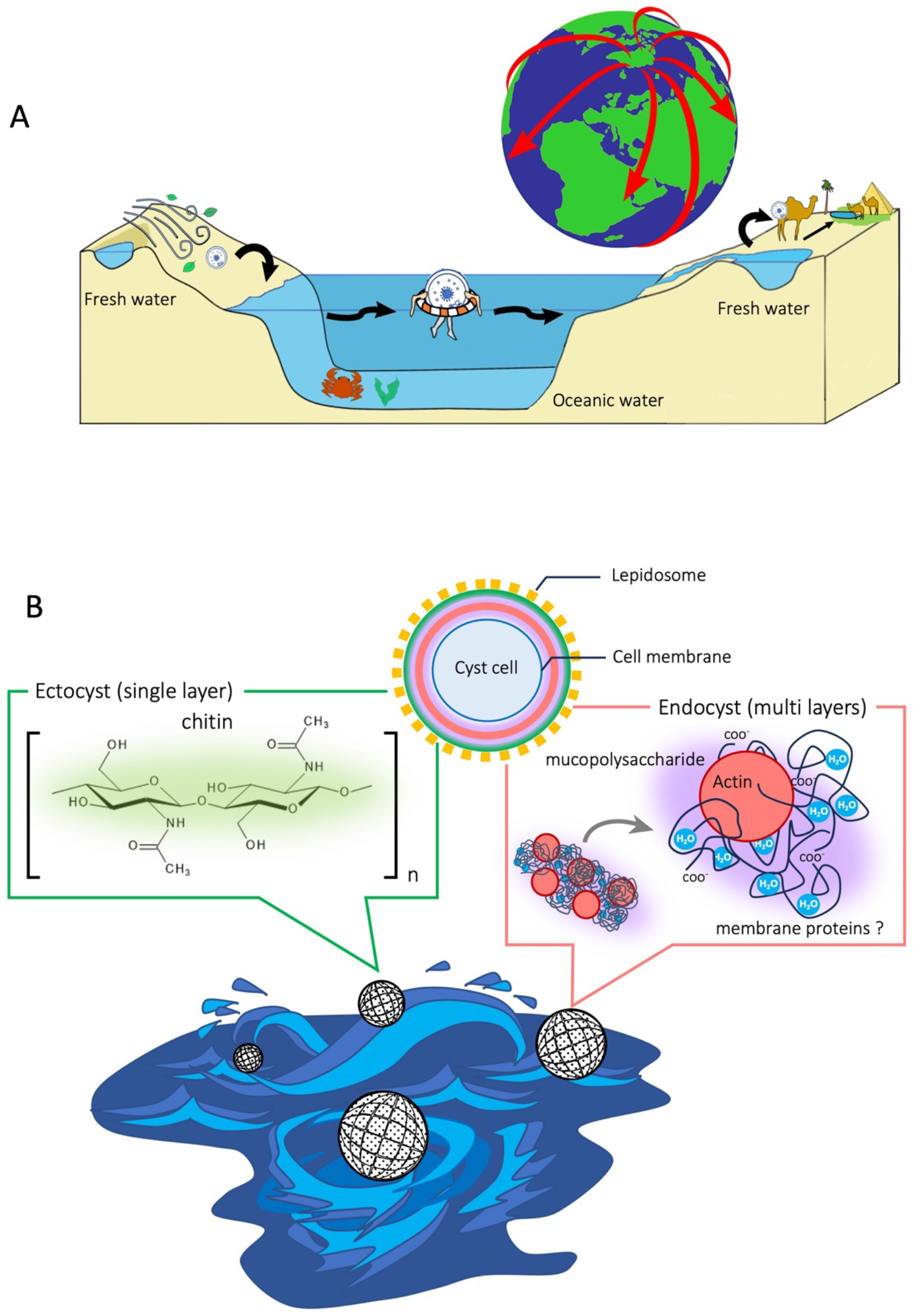
Illustration of our proposed model for oceanic dispersal of freshwater protists (A) and possible protection from mechanical stress by the chitinous ectocyst (B). (A) Freshwater ciliates such as *Colpoda* live in a temporary aqueous environment and they form resting cysts when the water disappears due to desiccation. Cysts may be transferred by wind, animals or human activity. Cysts that have accidentally been transported to the ocean get a chance to travel. (B) During their ocean travel, the cysts face stormy seas and mechanical stresses; however, the chitinous ectocyst protects the cysts from the stresses.

The chitinous ectocyst protects against physical shocks including strong water currents and bubbles (Fig. 7B). In the ocean, this defense mechanism might work more effectively when the cells shrink to create a gap between the cell membrane and ectocyst, enhancing their ability to withstand external forces. Furthermore, the endocyst serves as an additional shield by absorbing moisture and preventing dehydration by the hypertonic sea water (Fig. 7B). Actin and some membrane proteins can form gel-like networks during phase separation (72). Upon reaching a new freshwater habitat, excystment occurs and the organisms begin to proliferate as *r-*strategists. In brackish water where proliferation is limited, they can still excyst and remain active, but encystation is suppressed as they seek a more favorable environment for proliferation. Overall, the resting cyst formation could contribute to global dispersal via oceanic transportation. It might be feasible to search for freshwater protists ‘sailing’ across the ocean through environmental DNA analysis; this is a topic for further research.

Protists, including ciliates, are ubiquitously abundant in both marine environments, with high osmotic pressure, and hypotonic, freshwater environments. These habitats are completely different and the species have evolved in parallel. Marine protists accumulate osmoregulatory solutes (73), while freshwater protists have evolved contractile vacuoles (74). Despite the parallel evolution, resting cyst formation could enable freshwater ciliates (*Colpoda*) to survive oceanic dispersal due to their salinity tolerance.

There is probably no other life force equivalent to the resting cysts of unicellular organisms. It has enabled survival in a variety of extreme environments during the history of the Earth and possibly even oceanic dispersal. One of the authors (YS) has been investigating the tolerances of cysts for over a decade, having been fascinated by their extraordinary viability. Even when the environment causes extermination of all sorts of life, protists are able to survive. Cyst formation is a most beneficial survival strategy enabling the continued existence of single cellular organisms. It continues to fascinate him and others.

## Acknowledgments

We are deeply grateful to Prof. Shigeki Fujiwara (Kochi University, Japan) for his great suggestion for this manuscript. We are grateful to Prof. S. Uchida and three students at NIT. Fukushima College, Japan Mr. Sena Kobayashi, Mr. Taiga Shimizu, and Mr. Taiki Ono for their kind supports for this study, and Dr. Makoto Ogata (Fukushima University, Japan), Dr. Noriko Yamauchi (Ibaraki University, Japan), for their great kind supports and suggestions.

## Author contributions

**R.S.** conducted experiments, analyzed data, contributed to discussion, and wrote manuscript. **H.Y., K.Y., T.Saito, S.H., R.K.**, **T.Sakai**, **M.G., A.O., H.N., Y.Saito, T.A.** conducted experiments, and analyzed data. **H.Y. and M.G.** also participated in writing manuscript. **T.Suzuki**, **T.T.** conducted experiments, analyzed data, contributed to discussion, and wrote manuscript. **M.W.** contributed to discussion and wrote manuscript. **K.K.**, **F.S.**, **K.W.** contributed to discussion and participated in writing manuscript. **Y.Sogame.** designed the research, contributed discussion, and wrote draft of the manuscript. All authors contributed to review and editing of the final version of manuscripts.

## Data and materials availability

The sequence reads have been deposited in the DDBJ Sequence Read Archive (DRA)/SRA under the accession numbers DRR275117-DRR275118. All data are present in the paper and/or the Supplementary Materials.

