## Supplemental legends, Fig. SM1, Table SM1, SM2, SM3 for "Can *Colpoda* travel across oceans? Salinity tolerance of resting cysts may enable global dispersal of the species"

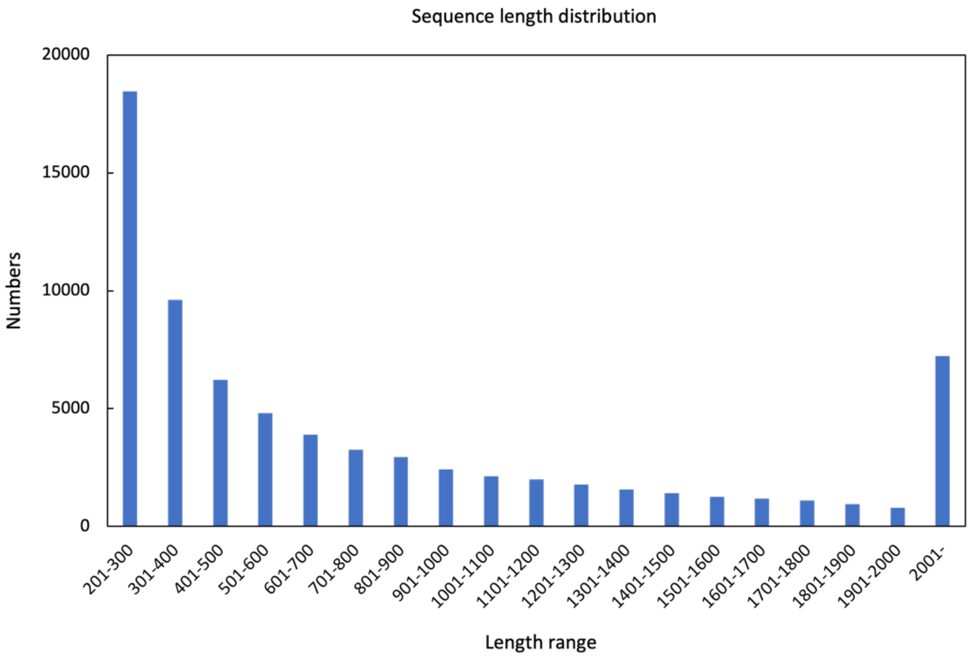


**Fig. SM 1.** Distribution of unigenes length from transcriptomic analysis of *C. cucullus* vegetative cell and cysts.

| Sample | Clean reads |
|---|---|
| Vegetative | 13,518,200 |
| Cyst | 12,733,840 |

**Table SM1.** Sequence results

| Total sequences | Total bases | Max sequence length (bp) | Min sequence length (bp) | Average length (bp) | Median sequence length (bp) | N50 length (bp) | (G+C)s | (A+T)s |
|---|---|---|---|---|---|---|---|---|
| 73,011 | 65,639,138 | 17,589 | 201 | 899.03 | 544 | 1,443 | 39.59 | 60.41 |

**Table SM2.** *De novo* sequence assembly

| Gene id | Gene name | Primer F | Primer R |
|---|---|---|---|
| Vegetative cell specific genes | | | |
| TRINITY_DN26936_c0_g2_i1 | V-type proton ATPase 16 kDa proteolipid subunit | AATGTAGCGCGTTGTTTGCC | ATTTTCGCCGAAGCTCTTGG |
| TRINITY_DN24844_c0_g1_i1 | Ubiquitin-conjugating enzyme E2-16 kDa | AATTCCACTGGCAAGCAACC | TGGGGTGATAGATTCTGGTGAC |
| TRINITY_DN12541_c0_g1_i1 | Eukaryotic translation initiation factor 5A-3 | ATCTTTTCTTGGCCGACAGC | TGGCATTAAGGCTGCTTTCG |
| TRINITY_DN29395_c1_g1_i1 | Cell division control protein 48 homolog A | ATATGCCGCCAGACAAAGTG | TTGTTGTTGACCACCGCTTG |
| TRINITY_DN39536_c0_g1_i1 | Cathepsin D | AAAAGCCAGACTGCACCAAC | AACACCCAAAAGGCATTGGC |
| Cyst specific genes | | | |
| TRINITY_DN29333_c1_g2_i1 | Calcium-dependent protein kinase 1 | TGGTGGTTTTTGCCTTCACG | ACAACCAGCCAACACAAGAC |
| TRINITY_DN34155_c0_g1_i1 | Growth arrest-specific protein 8 homolog | TTGGTCAATGAGCTCCAGAGAG | TTGGCACCAAAGGATCGAAC |
| TRINITY_DN33299_c0_g1_i1 | Parkin coregulated gene protein homolog | TTTGGGGGCTTGGTTTGTTC | AGCGAAACCGGAATCCAAAC |
| TRINITY_DN21381_c0_g1_i1 | ADP-ribosylation factor-like protein 3 | AGCGCCATCATCAAATCCTG | ACACAACTGATCACGGAAGC |
| TRINITY_DN480_c0_g1_i1 | Zinc finger protein 143 | ATGCAAACACGCAGGTTGTG | TTGGCACACAAAAGGTCTGG |
| Internal control genes | | | |
| TRINITY_DN22777_c0_g1 | Transcription elongation factor SPT4 | TCCTTCGAAGCTTGGTGTTG | ATGCCTTACATGCCGTCTTG |
| GenBank, accession No. X94348.1 | Colpoda sp. gene encoding for alpha-tubulin, partial | CGGTAAGGAAGATGCTGCCA | GGGACGAGGTTGGTTTGGAA |
| CHS5 gene | | | |
| TRINITY_DN26596_c0_g1 | RecName: Full=Cell fusion protein cfr1; AltName: Full=CHS5-related protein 1 | ACAAAACCGCAACTCCATCC | ACAAAACCGCAACTCCATCC |

Table SM3. List of gene id, gene name, and primers.

Table SM4. All transcriptome data in this study.

Table SM5. Results of gene enrichment analysis. Yellow color enhanced GO. terms were significant differences between vegetative cells and cysts. The positive number log2 transformed fold change represent upregulation in vegetative cells, while the negative represents upregulation of cysts.
